# Porosome reconstitution reverses Alzheimers in human brain organoids

**DOI:** 10.1101/2025.11.13.688334

**Authors:** Bhanu P. Jena, Won Jin Cho, Douglas J. Taatjes, Sushmita R. Patil

## Abstract

A central challenge in overcoming diseases resulting from defects in the cellular nanoscale multiprotein porosome complexes, is the consequent alteration of other proteins within the complex. By exploiting the distinct porosomes present in different tissues, Porosome Therapeutics, Inc., has developed a clinically relevant approach to incorporate (reconstitute) functional wild type porosomes into the plasma membrane of diseased cells to overcome secretory defects. In a combinatorial strategy of reprogramming the neuronal secretory and metabolic components, we established a dual-target therapeutic framework that repairs synaptic and metabolic defects to counteract neurodegeneration in Alzheimer’s. Utilizing iPSC derived human brain organoids, as recommended by the NIH and FDA, we demonstrate both the morphological and functional reversal of Alzheimer’s following porosome-apigenin treatment. Using the UHD-CMOS-MEA System developed jointly by Sony Semiconductor Solutions (Sony), SCREEN Holdings (SCREEN) and VitroVo, we were able to monitor thousands of neuronal firing in the brain organoids, helping in assessing the restoration of neuronal function in Alzheimer’s following porosome-apigenin therapy.

## INTRODUCTION

Alzheimer’s disease (AD) is characterized by progressive neurodegeneration and cognitive decline, driven by disruptions in neuronal metabolism and secretion^1-4^. A central challenge in Alzheimer’s disease is understanding the mechanism of neuronal secretory dysfunction. In an earlier study^5^, by exploiting AI, we revealed how beta-amyloid disrupts key protein interactions within the neuronal secretory machinery -the porosome [Fig 1, previously published in-part, is presented for clarity] ^5-8^. Using postmortem hippocampal analysis and AD neuronal cultures, we demonstrated that there is a marked depletion of porosome proteins, including SNAP-25, Syntaxin-1A, and ATP1A3 [Fig 2, previously published in-part, is presented for clarity], establishing a mechanistic link between Aβ toxicity and synaptic failure^9^. Furthermore, a major challenge in treating diseases resulting from defects in the multiprotein porosome complexes, is that the alteration of one protein within the complex impacts other proteins. Hence the incorporation of the entire functional porosome nanomachine into the plasma membrane of the diseased cell becomes necessary [Fig 3]. As a result, porosome reconstitution into AD neurons restores the missing proteins in AD neuronal cultures, reduces oxidative stress and cell death^9^.

**Figure 1:**
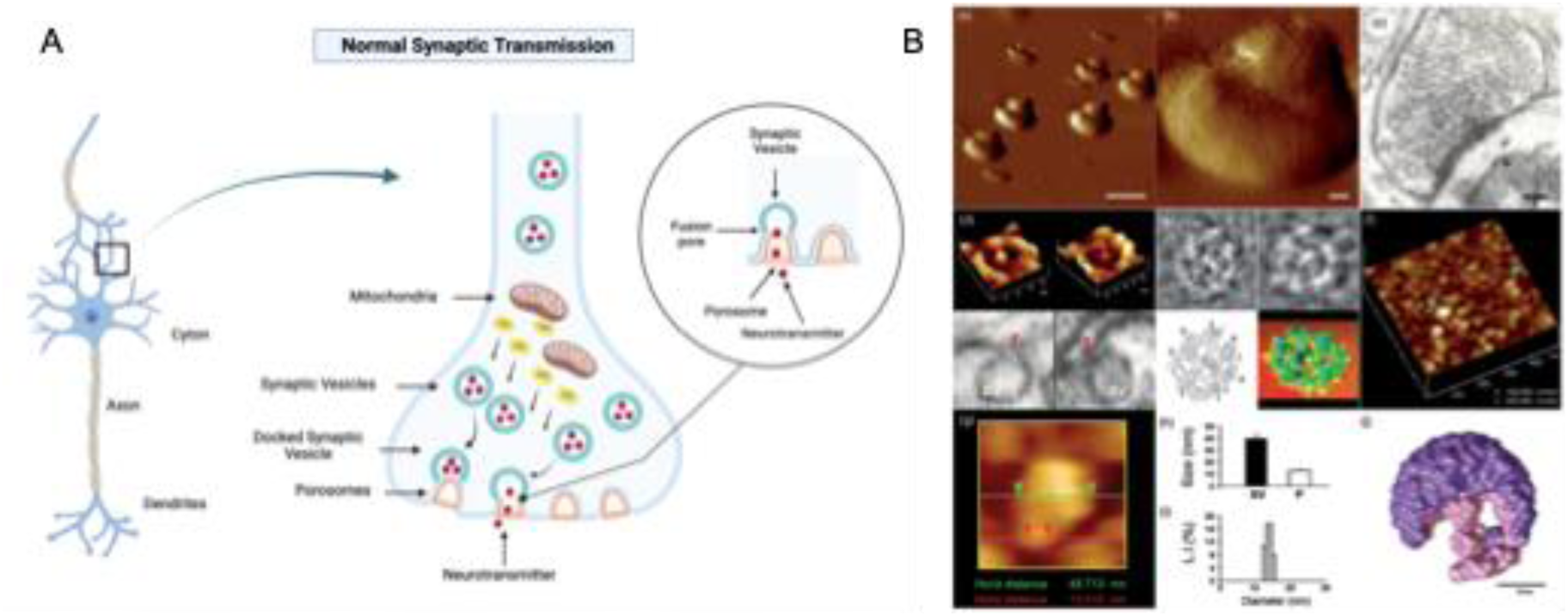
Fifteen nanometer cup-shaped neuronal porosome secretory nanomachines at the presynaptic plasma membrane of nerve terminals are required for neurotransmission. **(A)** Schematic illustration of a synapse at the nerve terminal in neurons, demonstrating normal mitochondrial function, energy (ATP) generation, and porosome-mediated neurosecretion. **(B)** Native neuronal porosome structure and organization. (a) Low- (Scale =1 μm) and (b) high-resolution (Scale 100 nm) atomic force microscope (AFM) micrographs of isolated rat-brain nerve endings or synaptosomes in buffer solution. (c) Electron microscope (EM) picture of a nerve ending (Scale = 100 nm). (d) Native neuronal porosome complex at the presynaptic membrane (Figure d top left), and of an isolated porosome complex reconstituted into lipid membrane (Figure d top right). Note the native and reconstituted porosomes are morphologically identical. Lower panels are two EM micrographs demonstrating synaptic vesicles (SV) docked at the base of a cup-shaped neuronal porosome having a central plug (red arrowhead). (e) EM, electron density, and 3D contour maps provide a nanometer-scale resolution of protein assembly within the neuronal porosome complex. (f) The cytosolic compartment of an isolated nerve ending or synaptosome demonstrating SV (blue arrowhead) docked at the base of porosomes (red arrowhead). (g) A SV docked at a porosome. (h) Measurements (n = 15) using AFM of SV (SV, 40.15 ± 3.14 nm) and porosomes (P, 13.05 ± 0.91 nm) at the presynaptic membrane. (i) Photon correlation spectroscopy performed on millions of isolated neuronal porosomes suspension in physiological buffer solution measure on average around 15 nm with a distribution of 12–17 nm. (j) X-ray solution scattering (SAXS) of the averaged 3-D structure of SV (purple) docked at the base of the native neuronal porosome complex (pink). SV–porosome complex using EM, AFM, and SAXS demonstrate similar morphology^5-8^.

**Figure 2:**
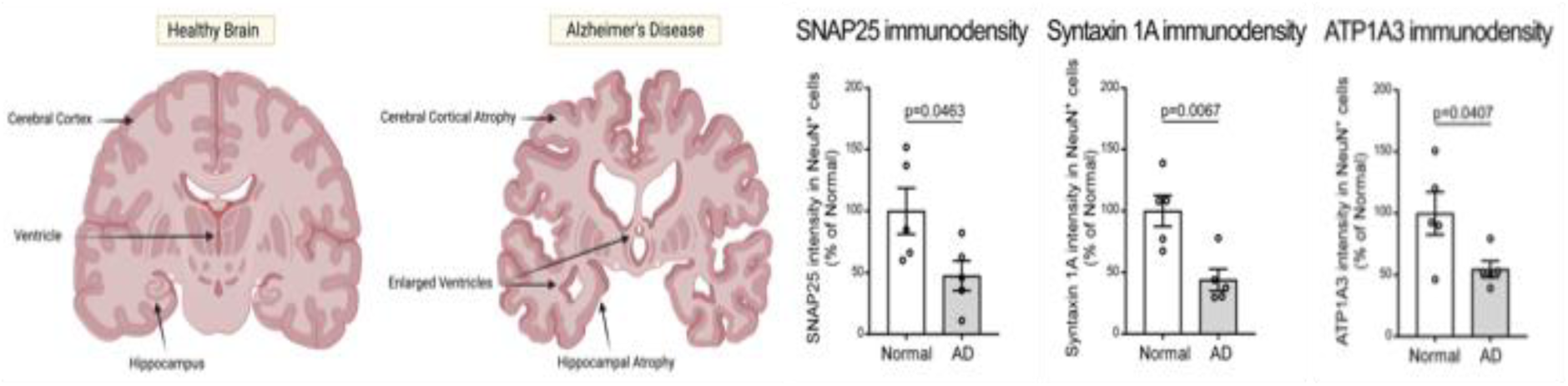
Immunoblot analysis of the hippocampus of AD patient postmortem brains demonstrate depletion of porosome proteins SNAP-25, Syntaxin-1A, and ATP1A3, as we previously report^9^. Postmortem hippocampus sections from control subjects and AD patients (n = 5 individuals/group) were stained with antibodies of anti-SNAP25, anti-Syntaxin 1A and anti-ATP1A3. The intensity of SNAP25, Syntaxin 1A, and ATP1A3 labeling was quantified and shown in bar graphs, respectively. All data are presented as mean ± SEM. Statistical significance was determined by unpaired Student’s t-test^9^.

**Figure 3:**
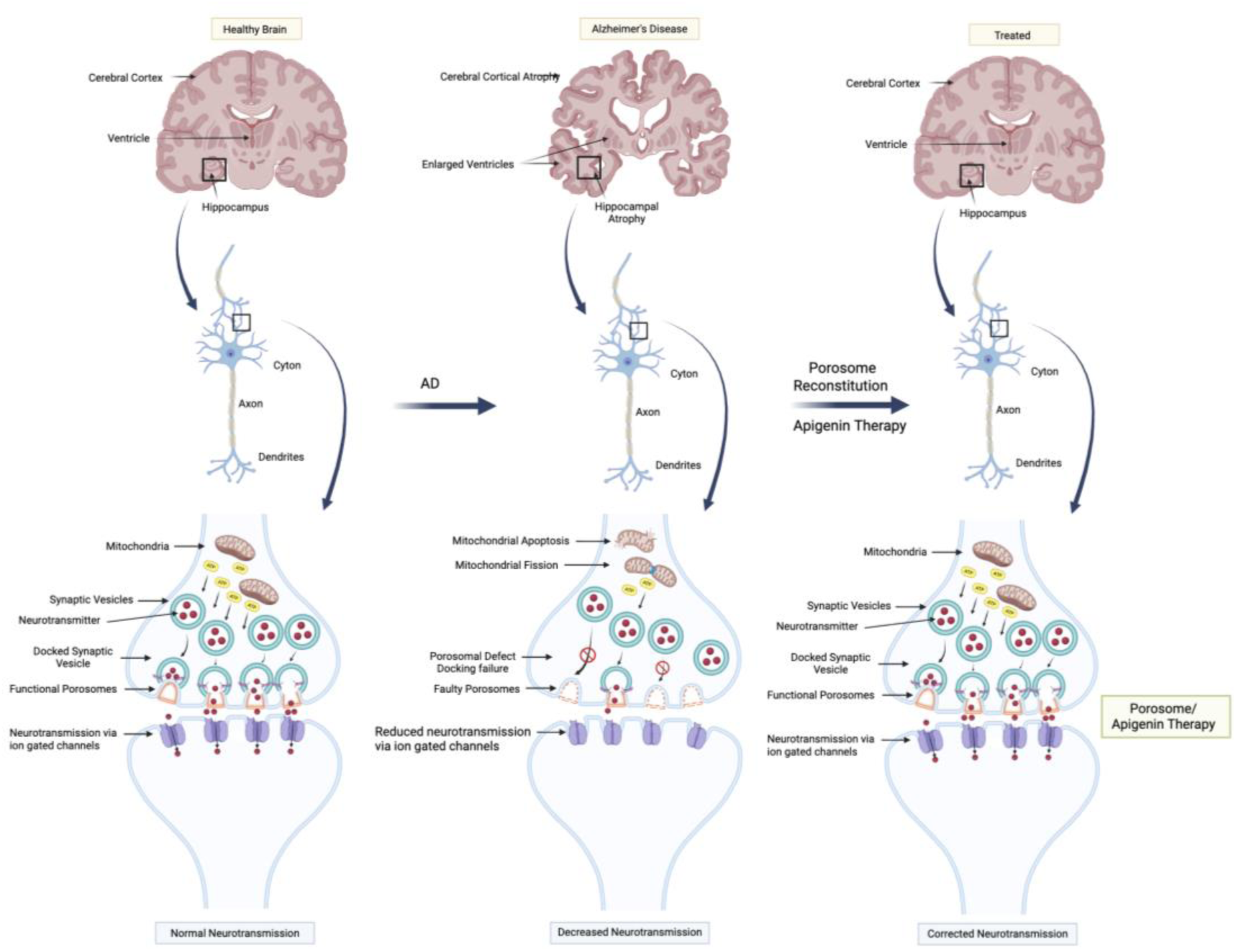
Schematic illustration of how porosome reconstitution therapy may overcome defects in neurotransmitter release in Alzheimer’s and other neurological diseases resulting from neurotransmitter release defects.

In the current study, a combinatorial strategy of reprogramming the neuronal secretory machinery (porosome reconstitution) and metabolic components (using the small molecule bioflavonoid Apigenin: 4′,5,7-trihydroxyflavone), we established a dual-target therapeutic framework that repairs synaptic and metabolic defects to counteract neurodegeneration in Alzheimer’s and potentially for other neurosecretory disorders. Utilizing iPSC derived human brain organoids, as recommended by the NIH and FDA, to assess the treatment of neurological secretory disorders such as those in Alzheimer’s, we demonstrate the morphological and functional reversal of Alzheimer’s in human AD brain organoids following porosome-apigenin therapy. Using an ultra-high-density CMOS-based microelectrode array (UHD-CMOS-MEA) as previously described^10,11^ we demonstrate the restoration of normal neuronal function in brain organoids with Alzheimer’s following therapy.

## MATERIALS AND METHODS

### Porosome Isolation and Reconstitution

Human neuronal cell line SH-SY5Y were used to isolate porosomes for reconstitution. RIPA buffer (10 mM Tris-HCl, pH 8.0 • 1mM EDTA • 0.5mM EGTA • 1% Triton X-100 • 0.1% Sodium Deoxycholate • 0.1% SDS • 140mM NaCl • Dilute with dH_2_O) containing 0.1mM PMSF, 1mM ATP, and a cocktail of protease inhibitors, was used to lyse the cells. Protein in all fractions was estimated using BCA Protein assay Kit (Thermo Fisher Cat. No. 23227, Rockford, IL 61101, USA). A total of 500 μg of the supernatant proteins were incubated with 2 μg of human monoclonal antibody raised against SNAP-25 overnight, followed by incubation with 40 μl of 50% protein AG Magnetic agarose beads (Thermo Fisher., Cat. No. 78610) slurry for 1 h. The beads were washed with the binding/wash buffer (PBS with 150 mM NaCl) for 5 min at 4 °C with gentle agitation twice, then washed with deionized water once. The proteins were eluted with 100 uL of 0.1 M glycine, pH 2.2 and neutralized with 25uL of 1M phosphate buffer, pH 7.5. Porosome reconstitution into human brain organoids in culture was achieved by exposing 0.125 μg/200 μL porosomes isolated from SH-SY5Y cells.

### Organoid Culture and Treatment

Human brain organoids (CIPO-BWL001K-20 Organoid) were purchased from Acro Biosystems (Newark, DE, 19711). Organoids were seeded in a 96-well Ultra-Low Attachment Plate with one organoid in each well containing 200 μL of cerebral organoid maintenance and differentiation medium. After 30 days, the experimental organoids were cultured with Aβ_1-42_ (10 μM,) to induce Alzheimer’s disease conditions. A minimum of seven days after incubation with Aβ_1-42_, organoids were either treated once with vehicle (AD) or with isolated human neuronal porosome (175 ng/well) (AD + porosome), apigenin (10μM) (AD + apigenin). The experimental organoids were continually exposed to 10 μM Aβ_1-42_. Organoid size was monitored every 24h. Organoids were maintained at 37 °C and 5% CO_2_. Apigenin (S2262) was purchased from Selleck Chemicals LLC (Houston, TX, USA).

### Preparation of oligomeric Aβ_1–42_

The Aβ_1–42_ (GenicBio Limited) peptides were dissolved in 1,1,1,3,3,3-hexafluoro-2-propanol (HFIP; 105228, Sigma-Aldrich) to a final concentration of 5 mM and placed in a chemical hood overnight. The next day, HFIP was further evaporated using a SpeedVac concentrator for 1 h. Monomer Aβ (5 mM) was prepared by dissolving Aβ peptide in anhydrous dimethyl sulfoxide (Sigma-Aldrich). The oligomeric Aβ peptides were prepared by diluting the monomer Aβ solution in Dulbecco’s Modified Eagle Medium (DMEM)/F12 and then incubating at 4 °C for 24 h.

### Confocal Immunofluorescence Microscopy

Organoids were fixed with 4% paraformaldehyde and stored prior to processing. Following fixation organoids were rinsed with PBS and embedded in 2% sea prep agarose to serve as a support matrix. Agarose blocks were hardened by fixation with 4% paraformaldehyde and embedded in paraffin. The paraffin blocks were then sectioned at 5 um thickness on a microtome and retrieved onto glass slides. Next, sections were subjected to antigen retrieval with DAKO Target Retrival solution at 100°C in a decloaking chamber, followed by blocking with 10% normal goat serum in PBS containing 0.1% triton X. The sections were then incubated with primary antibodies (rabbit anti-ATP1A3; Proteintech 10868-1-AP, 1:100 dilution or mouse anti-syntaxin 1; Santa Cruz sc-12736 1:50 diltuion) overnight at room temperature. Following rises with buffer the sections were incubated with secondary antibodies goat anti-rabbit (or mouse) conjugated to Alexa 555 (1:500 dilution) for 1 hr at room temperature. Following buffer rinses, the nuclei were counterstained with DAPI (1:500; Sigma-Aldrich D9542), and then the sections were imaged on a Nikon A1R point scanning confocal microscope.

### Image Analysis and Qualification

Immunocytochemistry (ICC) images were analyzed using *ImageJ* (National Institutes of Health, USA). Images were first converted to 8-bit grayscale and calibrated to scale using the microscope’s scale bar (µm/pixel). Nuclei were isolated by applying threshold segmentation to generate binary masks, which were refined using the “Watershed” function to separate closely spaced nuclei. To ensure spatial consistency across samples, multiple non-overlapping regions of interest (ROIs) measuring 100 × 100 µm² were defined across each image (typically six to seven per field) using the ROI Manager. Within each ROI, nuclei were analyzed using the “Analyze Particles” function to obtain morphometric parameters including area, perimeter, circularity, and centroid coordinates. The number of nuclei within each ROI was used to calculate nuclear density (nuclei per 100 µm²), and inter-nuclear distances were measured between nuclear centroids using the “Measure” tool. All measurements were exported as CSV files for further analysis. Data from multiple ROIs were averaged for each image, and results were expressed as mean ± standard error of the mean (SEM).

### Ultra-high-density CMOS-MEA neuronal recording

Brain organoids were recorded using an ultra-high-density CMOS-based microelectrode array (UHD-CMOS-MEA) as previously described^10,11^. Briefly, organoids were placed directly on the PEI-coated electrode surface and maintained at 37 °C and 5% CO₂ during recording. Field potential imaging (FPI) signals were sampled at 2–10 kHz, and data preprocessing and propagation analyses were performed following the same procedures described in Yokoi et al^11^.

## RESULTS

### Porosome reconstitution rescues compromised growth of iPSC derived human brain organoids with AD

While transgenic mice engineered with a combination of mutations in *APP* and other AD-associated genes, and or by overexpressing different apoE variants, have served as informative models for AD, particularly in assessment of pathological and behavioral features, these transgenic mouse models have major limitations^8^. Extensive neuronal loss for example, has not been observed in such AD mouse models, as opposed to in human AD patients. However, in recent years, with the rapid development of iPSC derived engineered human brain organoid 3D systems and their reprogramming to generate AD phenotype by exposing them to Aβ_(1-42)_ amyloid peptide, has become a well-established AD model to identify effective therapies for the disease^12^. Alzheimer’s disease (AD) pathology can be induced in iPSC-derived human brain organoids by direct exposure to exogenous beta-amyloid (Aβ) peptides or by using brain extracts from AD patients. This approach can mimic key AD features, including Aβ aggregation and tau phosphorylation, in control organoids.

Using established approaches, AD human brain organoids were generated and used to test the potency and efficacy of porosome reconstitution therapy. Organoids were seeded in a 96-well Ultra-Low Attachment Plate with one organoid in each well containing 200 μL of cerebral organoid maintenance and differentiation medium. After 30 days, the experimental organoids were cultured with Aβ_1-42_ (10 μM,) to induce Alzheimer’s disease conditions. Seven days after incubation with Aβ_1-42_, organoids were either treated once with vehicle (AD) or with isolated human neuronal porosome (175 ng/well) (AD + porosome), apigenin (10μM) (AD + apigenin). The experimental organoids were continually exposed to 10 μM Aβ_1-42_. Organoid size was monitored every 24h. Results from the study [Fig. 4] show that porosome reconstitution and apigenin treatment overcomes the negative impact of beta amyloid (Aβ_1-42_) on the growth of human brain organoids in culture. There is over a 50% rescue following just 15 days after porosome treatment [Fig. 4].

**Figure 4:**
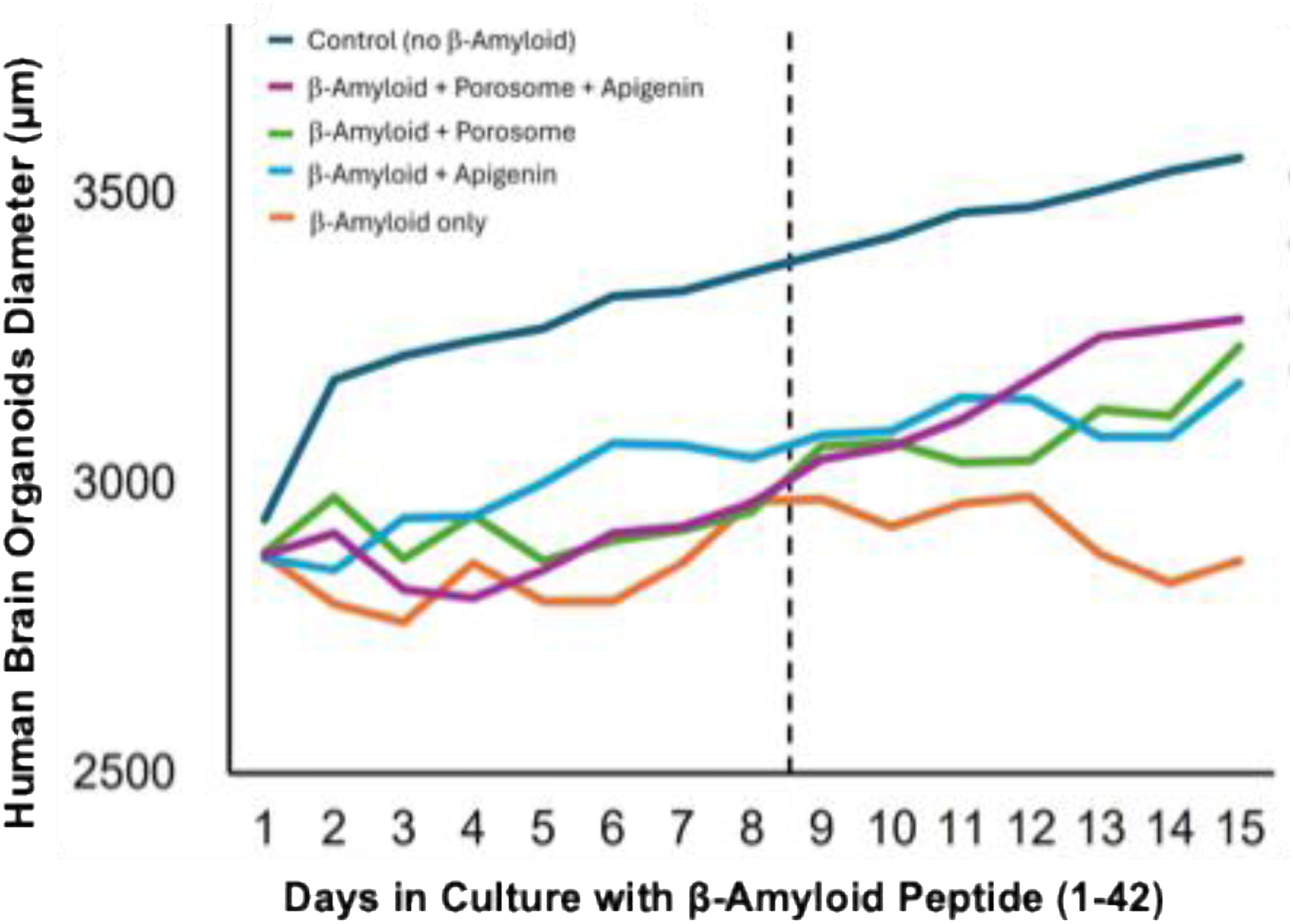
Porosome reconstitution and apigenin treatment overcomes the negative impact of beta amyloid (Aβ_1-42_) on the growth of human brain organoids in culture. There is over a 50% rescue following just 15 days after porosome treatment.

### Normal cellular morphology is restored by porosome reconstitution in AD organoids

To determine the impact of the porosome therapy on the cellular morphology in human brain organoids with AD, control, AD, and AD organoids treated once with porosome-apigenin were fixed and processed and immune stained using antibody to the porosome protein ATP1A3 gene product (the alpha 3 subunit of the sodium-potassium pump), and the nuclear stain DAPI. Results from this study reveals that in control brain organoids, there is a near uniform distribution of the ATP1A3 immunoreactivity (red) and the DAPI blue nuclear stain [Fig 5]. However, in AD organoids, there appears major gaps (seems devoid of tissue) within the organoid and a clustering of porosomes. The nuclear blue staining DAPI, stains poorly, there is a reduction of nucleus number, size, and inter-nuclear distance. However, following one treatment of porosome-apigenin, a near uniform distribution of the ATP1A3 immunoreactivity and the DAPI blue nuclear staining is restored to control levels. Consequently, the nucleus number, size, and inter-nuclear distance are also restored to normal levels [Fig 5].

**Figure 5:**
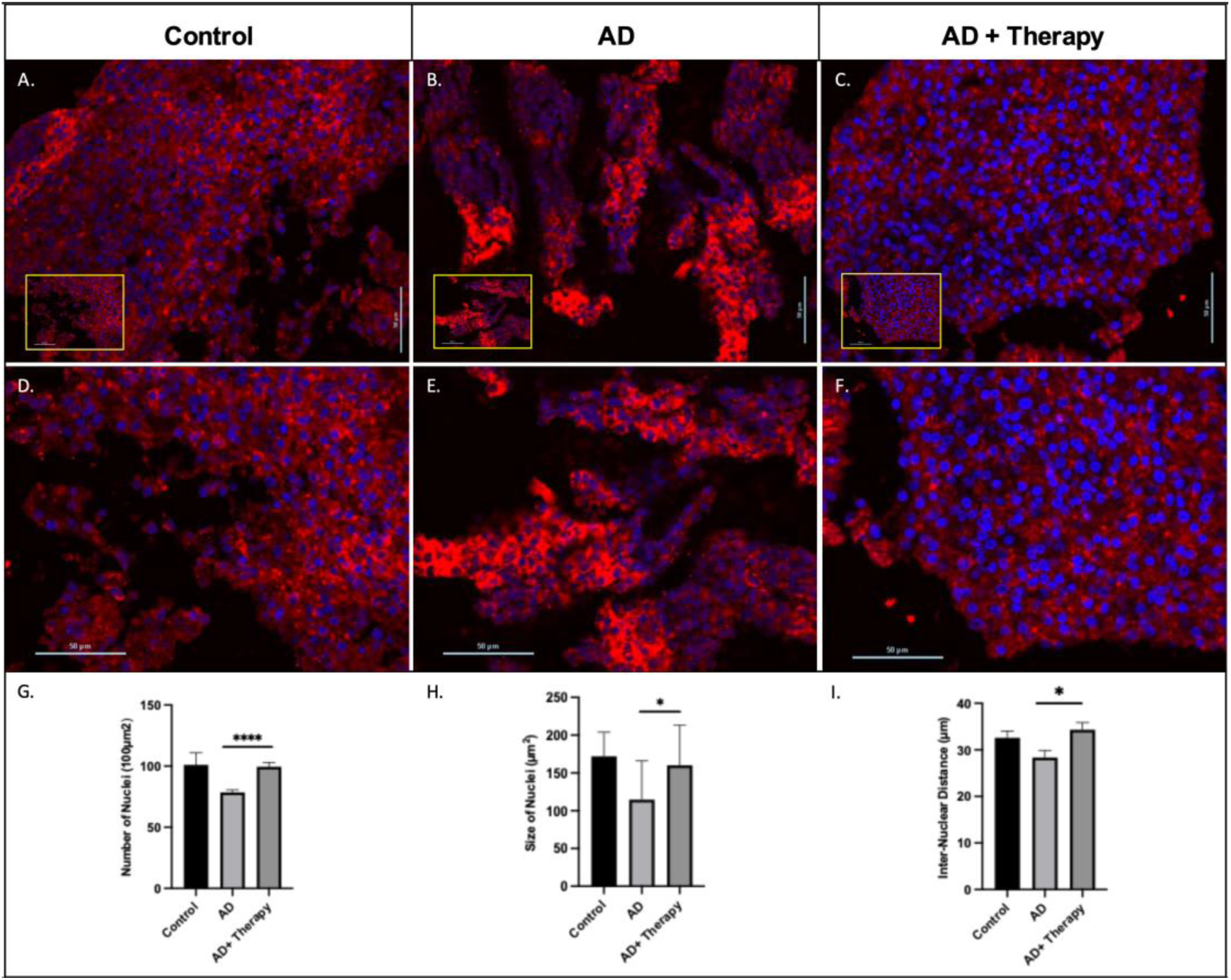
Porosome-apigenin treatment restores the morphology of AD neurons to normalcy. Control (**A, D**) normal human brain organoids exhibit uniform distribution of the porosome protein ATP1A3 (red) and the DAPI blue nuclear stain. In contrast, neurons in the AD organoids (**B, E**) show loss of neurons and clustered ATP1A3 protein and loss of DAPI nuclear staining. Treatment of AD organoids with porosome-apigenin (**C, F**) reverses AD neuron morphology to normal control levels with neuron regeneration, uniform distribution of the ATP1A3 porosome protein and the return of nuclear staining with DAPI. Note that nuclear size (**H**), number (**I**) and inter-nuclear distance (**J**) significantly reduced in the AD organoids, return to control levels following porosome-apigenin treatment. Data from multiple ROIs were averaged for each image, and results were expressed as mean ± standard error of the mean (SEM).

### Porosome reconstitution in AD organoids restores neuronal function

To determine the long-term impact of the porosome therapy on neuronal function in mature AD human brain organoids, 100-day old AD organoids were treated once with porosome-apigenin [Fig 6]. Thirty days following this treatment, neuronal activity was evaluated using the ultra-high-density CMOS-based microelectrode array (UHD-CMOS-MEA) system. The chip with more than 230,000 microelectrodes and the schematic diagram of a brain on a chip is shown for clarity [Fig 7]. A comparison of the neuronal activity in control, AD and AD-treated organoids is provided in a screen shot [Fig 8] and firing rate assessed in each of them [Fig 9]. Just one treatment of porosome + apigenin to the AD organoids 30-days prior to testing, demonstrates the complete reversal and retention of neuronal function to control levels [Fig 9].

**Figure 6:**
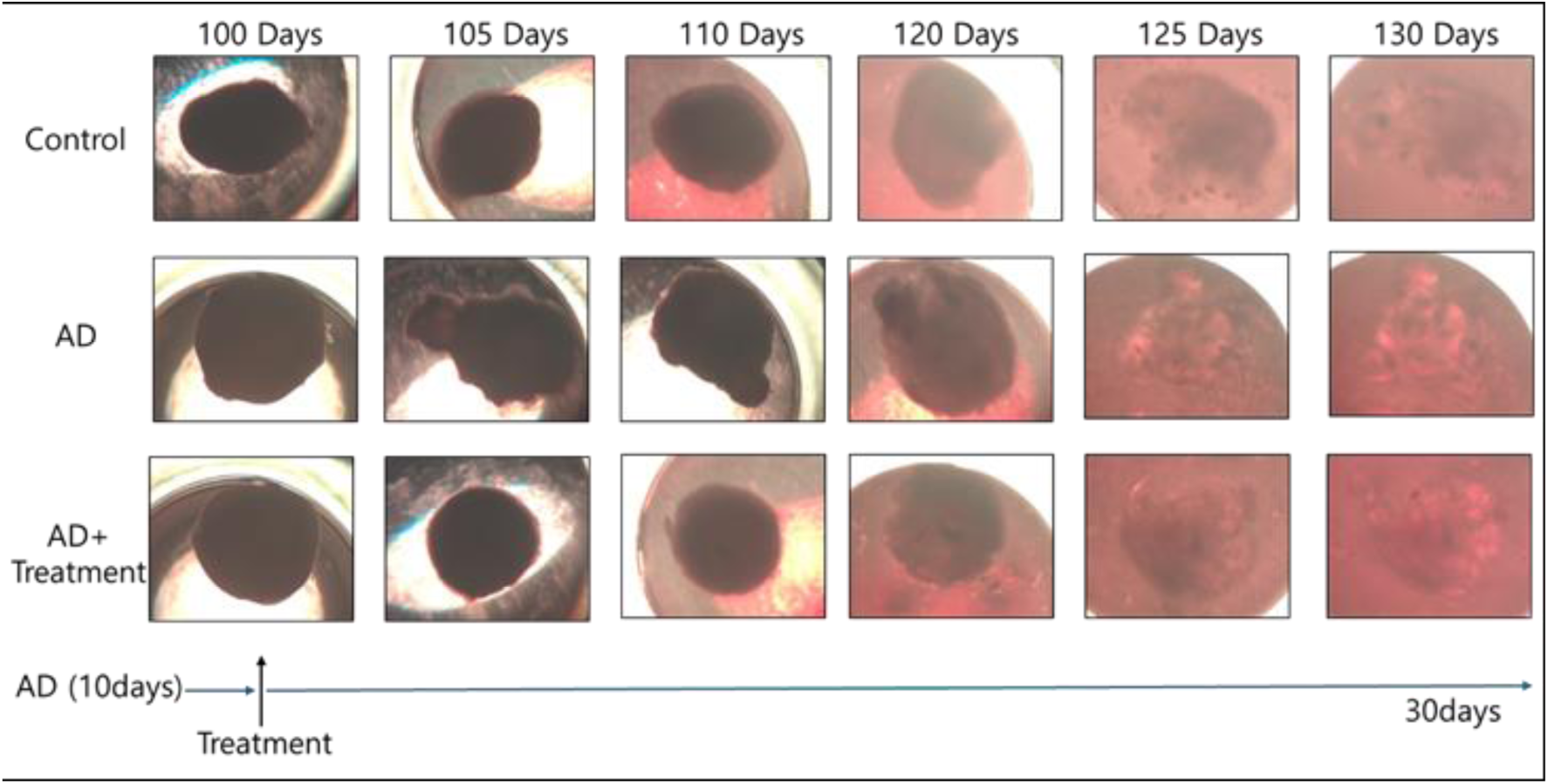
Light microscopy images of human brain organoids up to 130-days in culture in the presence or absence of beta amyloid (Aβ_1-42_ (AD) and AD organoids treated once with porosome + apigenin (AD + Treatment). Note the fragmentation if the AD organoid starting day 105, which is absent in the treated AD organoid.

**Figure 7:**
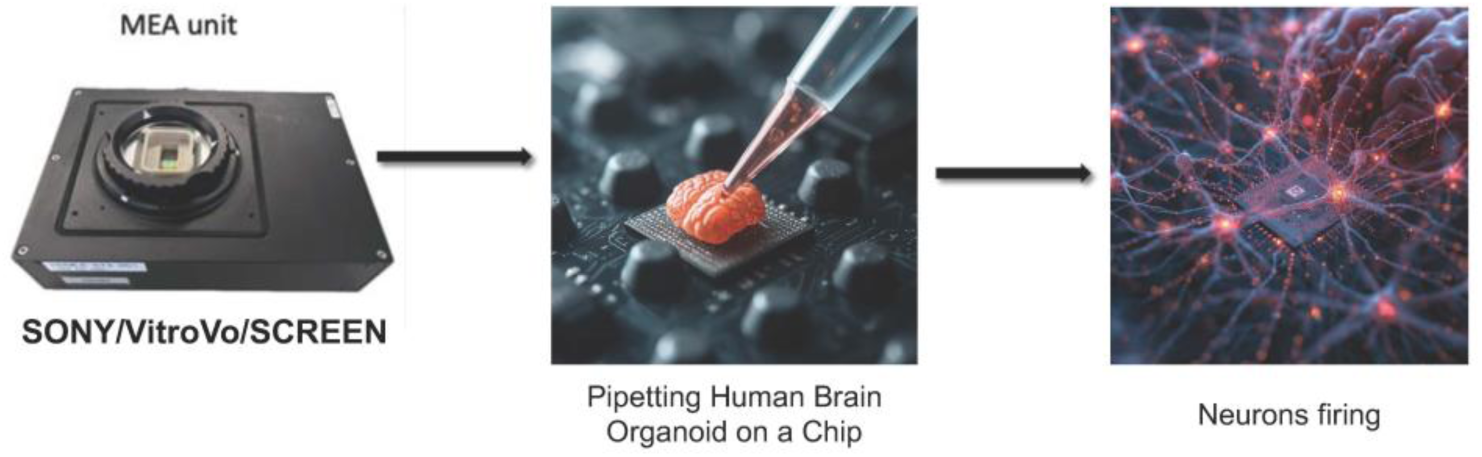
Ultra-high-density CMOS-based microelectrode array (MEA) chip and holding unit (far left) and schematics of ‘brain on a chip’ (center) and neurons firing (right).

**Figure 8:**
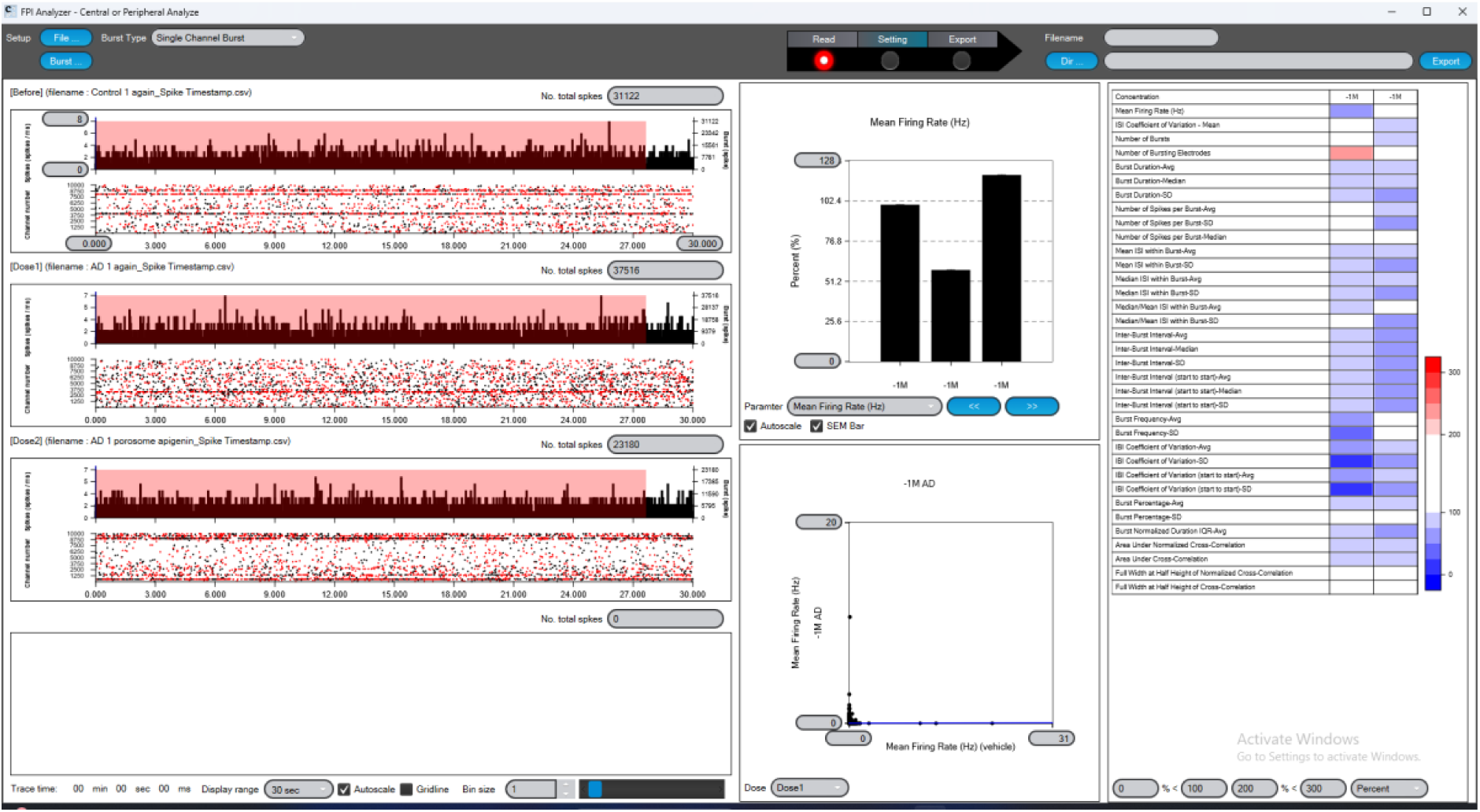
A screen shot depicting data acquisition on thousands of neuronal firing from three separate human brain organoid systems, namely from top-left ‘Control’, middle-left ‘AD’ and lower-left ‘AD + Treatment’ following 130-days in culture using the ultra-high-density CMOS-based MEA chip. Note to the top-center, the black bars reflecting the percent mean firing rate of neurons in the ‘Control organoids’ to the left (100%), in AD organoids (center) demonstrating a nearly 48% decrease, and ‘Porosome-Apigenin Treated organoids’ (right) showing a complete recovery to control levels.

**Figure 9:**
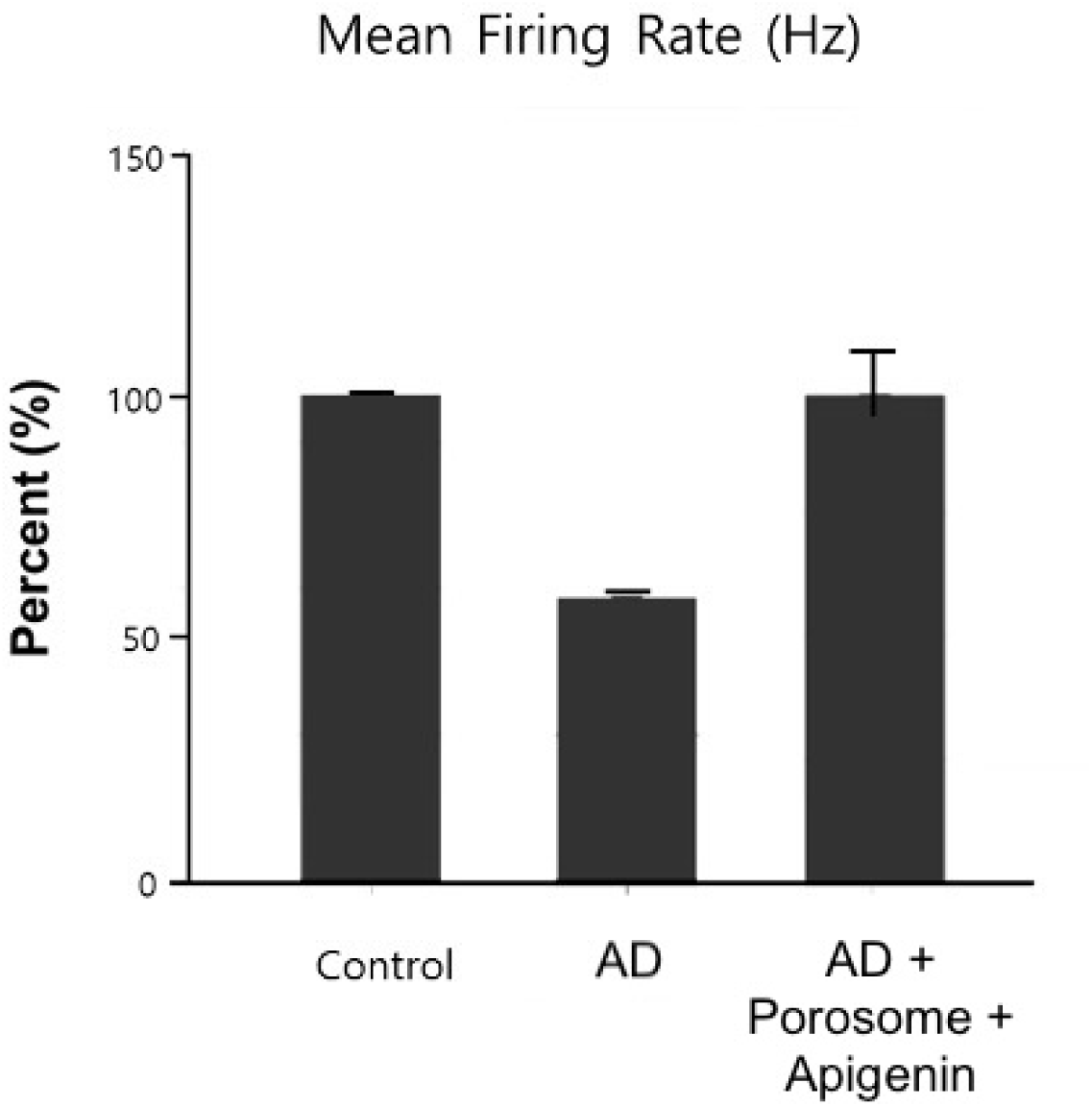
Poprosome treatment of AD brain organoids reverses their neuronal function to normal control levels. Data depicting the percent mean +/- SD of firing rate of neurons in human brain organoids following 130-days in culture using the ultra-high-density CMOS-based MEA chip. Firing rate in ‘Control’ organoids represent 100%, the AD organoids (β-amyloid-treated organoids) demonstrate a near 40% drop in the firing rate of neurons, and the reversal to control levels following porosome treatment (β-amyloid + Porosome + Apigenin). This recovery of AD organoid activity also reflects that after just one treatment of porosome + apigenin 30-days prior to the test, continues to maintain the complete recovery of neuronal function to control levels in AD organoids. Data represents 3-4 experiments with mean +/- SD of Control mean as100%, AD at 59.7% and AD + Treatment at 98.3.

**Figure 10:**
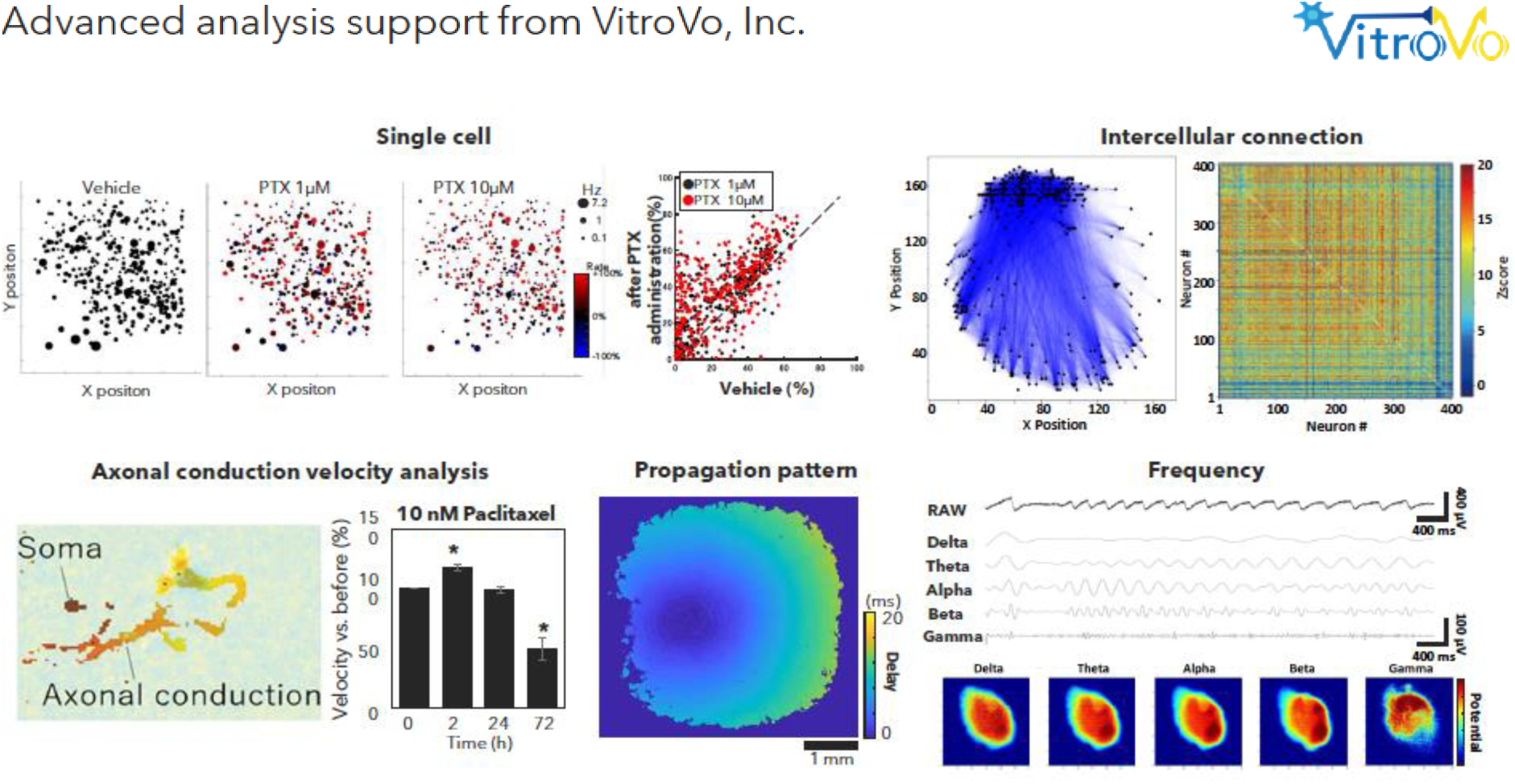
An example of the analysis by *VitroVo, Inc.* of data generated by the ultra-high-density CMOS-based MEA chip on brain organoids. Analysis of compound toxicity and efficacy using spike patterns, synaptic strength network activity, conduction velocity and frequency metrics, can all be assessed (*Courtesy of VitroVo, Inc.)*.

## DISCUSSION

This study lays the groundwork for porosome reconstitution therapy, a new and novel biologic that is capable of treating and reversing Alzheimer’s disease (AD). This therapy is a shift from conventional approaches focused primarily on amyloid clearance and inflammation. As previously reported^9^, despite diverse mechanisms of action, most therapeutic strategies have centered on amyloid clearance, neurotransmitter modulation, or inflammation control^13^. In contrast, the current study takes a systems-level approach to correcting both the metabolic and secretory defects in AD at every stage of the disease. In over 90% of AD patients, olfactory dysfunction precedes cognitive decline^14^, and the olfactory bulb of AD mice show a loss in expression of SNAP-25^15^. Since SNAP-25 levels in the cerebrospinal fluid (CSF) or in the blood of AD patients are significantly (p<0.0001) elevated, similar to the FDA-approved AD biomarker t-tau^16^, further supports that the porosome is negatively impacted very early on in the progression of the disease. Hence, administration of the proposed therapy could be via the nasal passage through the olfactory lobe, thereby bypassing the blood-brain barrier. Since the olfactory bulb is connected via nerves to the brain centers involved in learning, memory and emotion, the porosome or apigenin or both, could be deliverable through the nasal route. This is supported from our human brain organoid data, demonstrating that the addition of neuronal porosomes to the incubation medium holding the organoids is able to deliver the biologic deep into the organoid and its even distribution to the plasma membrane of neurons. Furthermore, even after one administration of porosomes, their presence and uniform distribution within the organoid is observed even 30-days following the treatment. This reflects both the efficacy, stability, viability of the porosome as a biologic, with no observable toxicity of the therapy. Using the large volume of data generated on spike patterns from the ultra-high-density CMOS-based MEA chip on brain organoids, we are currently obtaining using the analysis software developed by *VitroVo, Inc.* the compound toxicity and efficacy of porosome-reconstitution therapy. Using spike patterns, we are additionally obtaining the synaptic strength network activity, conduction velocity and frequency metrics, in AD brain organoids following therapy, laying the groundwork for clinical trials of porosome-apigenin therapy for Alzheimer’s.

## Contributions

Porosome-reconstitution therapy was developed at Porosome Therapeutics, Inc. The research design, interpretation and writing of the report was developed and performed by Bhanu P. Jena (B.P.J) and in-part conducted by Won Jin Cho (W-J.C) and Douglas J. Taatjes (D.J.T). W.J.C. and Yusuke Mori (Y.M) from SCREEN helped in conducting and recording the functional status of neurons in the organoids using the UHD-CMOS-MEA Chip. Morphometric assessment of immunostained organoid sections, and assembly of figures and the formatting of the manuscript were performed by Sushmita R. Patil (S.R.P). All authors participated in critical reading and discussion of the manuscript.

## Acknowledgments

The work presented in this article was supported by Porosome Therapeutics, Inc., and the Viron Molecular Medicine Institute, Boston, MA. Work presented in this article is patent protected by Porosome Therapeutics, Inc., Boston, MA. The functional status of neurons in the human brain organoids were assessed using the UHD-CMOS-MEA System developed jointly by Sony Semiconductor Solutions (Sony), SCREEN Holdings (SCREEN) and VitroVo. Juun Horie (J.H) from Sony, Bumpei Noda (B.N) from VitroVo, and Yusuke Mori (Y.M) from SCREEN organized the installation of the UHD-CMOS-MEA System in the Porosome Therapeutics, Inc., laboratory and organoid culture/recording support.

## Competing Financial Interest

This work is patent protected by Porosome Therapeutics Inc., W-JC and BPJ hold shares in Porosome Therapeutics, Inc.

